# Microstate Dynamics of Focused Attention Meditation

**DOI:** 10.64898/2026.01.19.700274

**Authors:** Chuong Ngo, Erkin Bek, Monika Stasytyte, Lionel Newman, Rodrigo Elizalde, Amit Kanthi, NK Manjunath, Christoph M. Michel

**Affiliations:** All Here SA, Geneva, Switzerland; Laboratory of Cognitive Neuroscience, Swiss Federal Institute of Technology Lausanne, Switzerland; Divison of Life Sciences, Swami Vivekananda Yoga Anusandhana Samsthana, Bengaluru, India; Department of Basic Neurosciences, University of Geneva, Switzerland

## Abstract

Focused-attention meditation provides a tractable model for examining how large-scale brain dynamics support attention and self-regulation. Using high-density EEG microstate analysis, we investigated how focused-attention meditation on the breath (Ānāpānasati) modulates intrinsic brain activity in 22 experienced practitioners, compared with baseline rest and deliberate mental imagery. Five canonical microstate classes (A-E) were identified. Meditation produced a robust reduction of Microstate C across coverage, duration, and occurrence, accompanied by increased presence of Microstates D and E (all Microstate x Condition interactions p < 0.0001). Source localization revealed that Microstate C was generated primarily in medial and lateral temporal regions including the hippocampus and parahippocampal cortex, whereas Microstate D involved posterior midline regions including the posterior cingulate cortex and precuneus, and Microstate E engaged frontoparietal and orbitolimbic networks. Together, these results indicate that focused-attention meditation reorganizes the temporal architecture of large-scale brain dynamics by downregulating microstate patterns associated with self-referential and memory-based processing while enhancing neural states supporting attentional stability and internal monitoring.

## 1. Introduction

Meditation offers a unique window into the dynamics of human consciousness, providing a stable and trainable framework for investigating how attention, self-awareness, and mental silence emerge from large-scale brain activity (Lutz et al., 2008; Dahl et al., 2015). While numerous neuroimaging studies have shown that meditation influences oscillatory rhythms, functional connectivity, and activity within the default mode network (DMN) (Fox et al., 2016), these approaches often lack the temporal precision needed to capture the rapid transitions that shape moment-to-moment experience. EEG microstates—transient, quasi-stable topographies lasting approximately 40–120 ms—provide an ideal method for tracking such fast-changing cognitive dynamics (Lehmann et al., 1987; Michel & Koenig, 2018). Conceptualized as the “atoms of thought,” microstates reflect fundamental building blocks of large-scale neural network activity and offers a dynamic tool to investigate underlying characteristics of human consciousness during rest (Custo et al., 2017; Zanesco 2024) or during altered states of consciousness (Bréchet & Michel 2022). Because meditative practices deliberately modulate *conscious states* — reducing self-related thoughts and mind-wandering while cultivating present-moment awareness (Fell et al., 2010; Lutz et al., 2015; Vago & Zeidan, 2016)— microstate analysis offers a powerful, mechanistically grounded approach for understanding how meditation reorganizes the temporal structure of conscious experience. Investigating meditation through the lens of microstates therefore not only illuminates the neural signatures of contemplative training but also contributes to broader models of how the brain organizes and stabilizes states of focused awareness, introspection, and mental stillness.

Building on this concept, a growing body of intervention research demonstrates that meditation training affects the brain’s intrinsic large-scale dynamics, as captured by EEG microstates. Zanesco et al. (2021) reported that two three-month Shamatha retreats led to significant decreases in microstate duration and strength across the canonical classes (A–D), along with altered transition probabilities, with subjective improvements in attentiveness and serenity showing strong explanatory power. Bréchet et al. (2021) observed a reconfiguration of microstate topographies (particularly microstates C and E) and their cortical sources, involving the right insular, the superior temporal gyrus, the superior parietal lobule, and the superior frontal gyrus bilaterally in participants after six weeks of app-based breath-focused training. Likewise, in an eight-week mindfulness-based stress reduction training (MBSR) training, Zarka et al. (2024) found significant reductions in the duration, occurrence, and coverage of microstate C, with microstate C parameters moderately correlated with the mindfulness facet of non-reactivity. Long-term style-specific expertise also shapes baseline EEG: Kopal et al. (2014) reported that practitioners of insight versus calm meditation exhibit distinct coherence patterns affecting the expression of microstates A–D even at rest. Together, these intervention studies indicate that meditation training reliably alters the spatio-temporal dynamics of large-scale brain networks.

Further evidence comes from studies measuring EEG microstates **during** meditation, showing how moment-to-moment contemplative experience is reflected in dynamic neural patterns. During open-monitoring (non-reactive attention) meditation, Zarka et al. (2021) reported a state-related decrease in microstates A and C and an increase in microstate B, with microstate temporal parameters also relating to mindfulness traits. In Transcendental Meditation, Faber et al. (2017) found significantly lower occurrence of microstates A and C during the “transcending” phase compared with undirected mentation. Early work by Faber et al. (2005) demonstrated that deep meditative absorption increases the duration of microstates B–D, while Panda et al. (2016) showed that meditation increased a DMN-linked microstate, with effects scaling with years of experience. More recent MEG work by D’Andrea et al. (2024) further revealed style-specific neural signatures, with focused-attention meditation increasing visual microstate MS1, whereas open-monitoring practice enhanced executive and ventral-attention microstates MS3 and MS5. Together, these studies suggest that meditation reliably modulates the presence of microstates associated with mind-wandering and attention depending on the specific meditation style.

Motivated by these prior observations, the present study examines microstate dynamics during Focused-Attention (FA) meditation in experienced practitioners. We conducted microstate analysis on high-density EEG recordings collected at Pyramid Valley International (Bengaluru, India), a large contemplative centre rooted in the Indian meditative culture whose primary practice is sustained awareness of the natural breath, known *as Ānāpānasati (“*Mindfulness of breathing”). *Ānāpānasati* is one of the most widely described classical breath-based meditation practices, appearing prominently in early Buddhist literature such as the Ānāpānasati Sutta (MN 118; Bodhi, 2015). Within modern contemplative science, mindfulness of breathing corresponds to the Focused Attention category of meditation practices (Lutz et al., 2008). More broadly, breath-focused attention is widely employed across diverse contemplative practices as a foundational method for cultivating attentional stability, functioning as a core or preparatory technique not only in Buddhist lineages (Buddhaghosa, 2011; Goenka, 1997) but also in Yogic concentration practices (Feuerstein, 2011), Zen breath-regulation training (Aitken, 1994), modern mindfulness programs including MBSR and MBCT (Kabat-Zinn, 1990; Segal et al., 2013), and various contemporary meditative approaches that use breath awareness as a gateway to deeper cognitive and experiential refinement (Brown et al., 2013).

In Ānāpānasati practice, attention is maintained on the tactile sensations of the breath— typically at the nostrils or along the natural flow of inhalation and exhalation—while minimizing cognitive elaboration, imagery, and conceptual evaluation. When distractions such as spontaneous thoughts, memories, or emotional reactions arise, practitioners gently acknowledge them and return attention to the breath, strengthening both sustained attention and meta-awareness (Dunne, 2015; Vago & Zeidan, 2016). Over time, this repeated cycle of focusing, distraction, and reorientation leads to increasing continuity of attention and a progressive quieting of self-referential and narrative mental activity, thereby reducing mind-wandering and enhancing cognitive stability. This core mechanism of attentional training has been well documented in contemplative cognitive research and is thought to support the development of sustained concentration and cognitive control through the systematic modulation of attentional networks (Lutz et al., 2008; Hasenkamp et al., 2012). In highly experienced meditators, sustained practice of Ānāpānasati may culminate in deep absorptive states traditionally described as Jhāna (Pāli), Dhyāna (Sanskrit), or Samādhi, characterized by markedly reduced internal dialogue, heightened perceptual clarity, and profound mental stillness (Wallace, 1999; Analayo, 2019). These states closely parallel descriptions of meditative absorption in the *Yoga Sūtra* of Patañjali (e.g., Yoga Sūtra I.2–I.3; Patañjali, trans. 1890), which emphasize the cessation of mental fluctuations and the stabilization of awareness in its own nature.

Although the underlying neuroscientific mechanism of FA meditation has been studied for decades, a recent systematic review (Lieberman et al., 2025) highlights substantial inconsistencies in existing literature, particularly in spectral analyses. The authors emphasize that conclusions based solely on electrode-level power measurements may be misleading, as scalp signals do not reliably reflect their underlying neural generators. They therefore recommend that future research incorporate spatially informed methods—such as EEG microstate analysis and source localization techniques including low-resolution electromagnetic tomography (LORETA)—to better characterize the neural mechanisms supporting FA.

In the present study, we follow this recommendation by applying microstate analysis to high-density EEG recorded during Ānāpānasati practice. We characterize the resulting microstate topographies during FA meditation, quantify their temporal properties (occurrence, duration, coverage), and reconstruct their cortical generators using LORETA. Finally, we assess how these microstate dynamics relate to mind-wandering, attentional stability, and self-awareness, thereby providing a more mechanistic account of the neural processes underlying FA meditation.

## 2. Methods

### 2.1 Participants and protocol

Twenty-five subjects participated in this study. Three subjects were excluded: two performed different meditation practices and one withdrew the consent after the recording. The remaining 22 subjects includes 8 male and 14 female. Subjects’ age ranged from 27 to 77 years (M = 50.6, SD = 15.4). They reported the following durations since beginning their meditation practice: 2–5 years (8 participants), 5–10 years (4 participants), 10–20 years (9 participants), and more than 20 years (4 participants). All participants practiced *Ānāpānasati* and were recruited through recommendation of the Pyramid Valley international. All subjects signed an informed consent form. The study was approved by the institutional Ethics committee of S-VYASA (RES/IEC-SVYASA/394/2025).

All participants followed the same fixed protocol. They were seated in a meditation room either on a chair or on the floor in meditation posture. The protocol consisted of three conditions, all performed with eyes closed: First, participants were instructed to sit still and relax for 3 minutes without meditating, allowing their mind to wander freely (Baseline). Second, they were instructed to visually imagine a scene from the past as vividly as possible for 3 minutes (Mental Imagery). Finally, participants performed their usual meditation practice for 20 minutes (Meditation). There was about 1-2 minutes break between the conditions.

### 2.2 Data acquisition and data analysis

EEG was recorded continuously using a 64-channel net with equidistant electrode positioning (ANT Neuro). Electrodes consisted of sponges soaked in saline water. Electrode impedances were maintained below 50 kΩ. Data were acquired with 5Z as the recording reference at a sampling rate of 500 Hz.

Preprocessing: EEG data were down-sampled to 250 Hz and band-pass filtered between 2-40 Hz using a zero-phase, 4th-order Butterworth filter. A 50 Hz notch filter was applied to remove line noise. The reference electrode was added as a zero-valued channel prior to re-referencing, and the data were subsequently re-referenced to the average reference. All recordings were visually inspected for artifacts. Bad electrodes were interpolated using spherical spline interpolation, and artifact periods were marked and excluded from further analysis. Finally, data were spatially smoothed using the spatial filter implemented in the Cartool software, as described in detail in Michel & Brunet (2019).

#### Microstate Segmentation

A modified k-means cluster analysis (Pascual-Marqui et al., 1995) was applied to the EEG of each subject across the three eyes-closed conditions (Baseline, Mental Imagery, and Meditation) using the open-source software Cartool, following the recommendations of Bagdasarov et al. (2024). The analysis consisted of two steps:

#### Individual-level clustering

The Global Field Power (GFP) peaks of each subject’s EEG across the three conditions were extracted and subjected to k-means clustering with 50 repetitions. The data were randomly split into epochs of 5 seconds covering 99.9% of the data. This resampling was repeated 45 times to produce more reliable clustering. For each epoch, the number of clusters was set to range from 4 to 12, and the optimal number for each epoch was selected using the meta-criterion—an aggregate measure of six independent criteria implemented in the Cartool software.

#### Group-level clustering

The optimal cluster maps from each of the 45 epochs of all subjects were then subjected to a second k-means clustering with 100 resampling trials and 100 repetitions. The optimal number of clusters was determined using the meta-criterion, which yielded 5 microstate classes.

#### Microstate Back-fitting

The optimal cluster maps resulting from the group-level clustering were back-fitted to each subject’s data across the three conditions. Using spatial correlation of GFP-normalized maps, each time point was labeled with the cluster map that showed the highest correlation (winner-takes-all labeling), with polarity ignored. Time points that showed correlations lower than 50% with any cluster map were not labeled. After back-fitting, temporal smoothing was applied (window half-size of 40 ms and Besag factor of 10; Pascual-Marqui et al., 1995). Segments shorter than 40 ms were removed by assigning the first half to the preceding segment and the second half to the following segment. The following parameters were extracted for each microstate class: mean duration (in milliseconds), coverage (percentage of time points labeled by the microstate), and occurrence (number of segments per second).

### Statistical Analysis

Statistical analysis was performed using GraphPad Prism. The three fitting parameters (duration, coverage, and occurrence) were subjected to two-way repeated measures ANOVA with microstate class and condition as within-subject factors. Post-hoc pairwise comparisons were performed using paired t-tests with correction for multiple comparisons by controlling the false discovery rate using the linear step-up procedure (Benjamini, Krieger, & Yekutieli (2006). Statistical significance was set at α = 0.05.

### Source Localization

After back-fitting, all time points labeled with a given microstate class were concatenated for each subject and each condition. For source localization, only time points with correlations >80% with the corresponding cluster map were retained. The 64 electrode positions were co-registered to the MNI-152 template head using a semi-automatic procedure implemented in Cartool. Sources were modeled using a 4-shell adaptive Local Spherical Model with Anatomical Constraints (LSMAC). This head model constructs local spheres with different radii for each electrode by estimating the thickness of the scalp, skull, cerebrospinal fluid, and brain tissue under each electrode (Brunet et al., 2011). As the source model, we used the Low-Resolution Brain Electromagnetic Tomography (LORETA; Pascual-Marqui et al., 1994) distributed linear inverse solution with 6926 solution points. The results were optimized using z-scoring implemented in Cartool, which eliminates activation biases (Michel & Brunet, 2019). The activity at each solution point was averaged across time points for each microstate and then averaged across participants. The group-level source distribution was thresholded to solution points above the 95th percentile of activation values, as done in previous work (Bagdasarov et al., 2022; Bréchet et al., 2019, 2021).

## 3. Results

### Microstate Segmentation

The meta-criterion applied to the group-level clustering identified 5 maps as the optimal number of clusters. These maps are illustrated in Figure 1. The topographies of the five maps were highly similar to the canonical maps reported in the meta-analysis of 50 studies by Tarailis et al. (2024) and the Meta-microstates from 40 studies reported by Keonig et al., (2024) and were labeled as microstate maps A through E.

**Figure 1:**
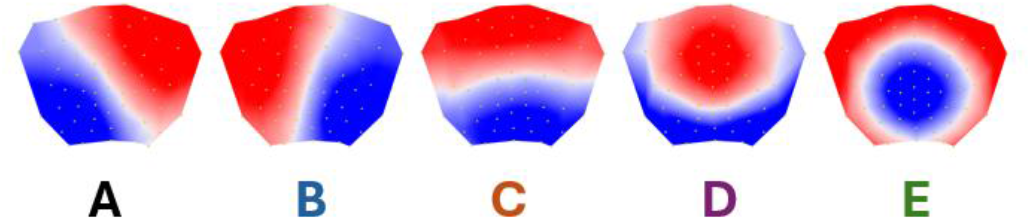
Group-level microstate segmentation identified five prototypical scalp topographies, labeled microstates A–E, as the optimal clustering solution based on a meta-criterion. The resulting maps closely resemble the canonical EEG microstate classes reported in large-scale normative studies and meta-analyses, including Koenig et al. (2024)

### Fitting Parameters

The fitting parameters (duration, coverage, and occurrence) were analyzed separately using two-way repeated measures ANOVAs with microstate class (5 levels) and condition (3 levels: Baseline, Mental Imagery, Meditation) as within-subject factors.

### Coverage

The two-way ANOVA revealed a significant main effect of Microstate (F(1.861, 39.08) = 32.05; p < 0.0001), and a significant Microstate × Condition interaction (F(2.560, 53.76) = 17.46; p < 0.0001). The main effect of Microstate reflected generally higher coverage of microstates A, B, and C compared to microstates D and E (Figure 2).

**Figure 2:**
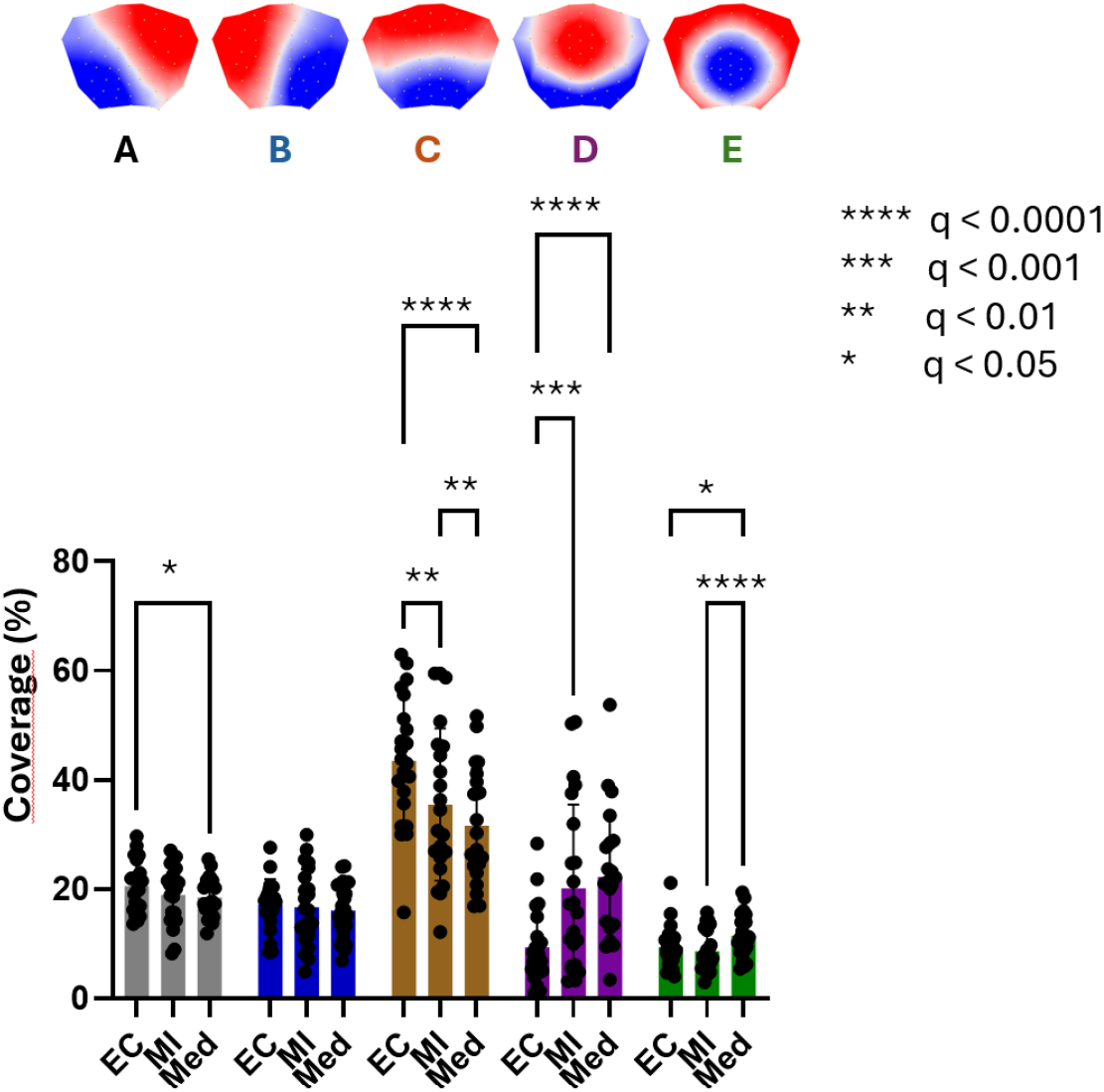
Microstate coverage across conditions. Mean coverage (%) of microstates A–E during Baseline (BL), Mental Imagery (MI), and Meditation (Med). A two-way repeated-measures ANOVA revealed a significant main effect of Microstate and a significant Microstate × Condition interaction. Overall, microstates A–C showed higher coverage than microstates D and E. Post-hoc comparisons (FDR-corrected) indicated reduced coverage of microstate C during MI and Med relative to BL, with a further reduction during Med compared to MI. In contrast, microstate D showed increased coverage during MI and Med relative to BL, with an additional increase during Med. Microstate E coverage increased during Med relative to both BL and MI, while microstate B showed a modest decrease during Med compared to BL. No significant condition effects were observed for microstate A. Significance levels are indicated in the figure.

Post-hoc pairwise comparisons corrected for false discovery rate revealed the following pattern for the Microstate × Condition interaction: Microstate C showed significantly decreased coverage during both Mental Imagery and Meditation compared to Baseline, with a further significant decrease during Meditation compared to Mental Imagery. Conversely, microstate D coverage significantly increased during both Mental Imagery and Meditation compared to Baseline, with an additional increase during Meditation compared to Mental Imagery. Microstate E coverage increased significantly during Meditation compared to both Baseline and Mental Imagery, while no difference was observed between Baseline and Mental Imagery. Microstate A showed a modest but significant decrease in coverage during Meditation compared to Baseline. Microstate B coverage did not differ significantly across conditions.

### Duration

The two-way repeated measures ANOVA revealed significant main effects for both Microstate (F(1.724, 36.20) = 31.21; p < 0.0001) and Condition (F(1.554, 32.63) = 5.721; p = 0.012), as well as a significant Microstate × Condition interaction (F(2.707, 56.84) = 16.16; p < 0.0001). Microstate C showed generally longer mean duration than the other microstates across all conditions (Figure 3).

**Figure 3:**
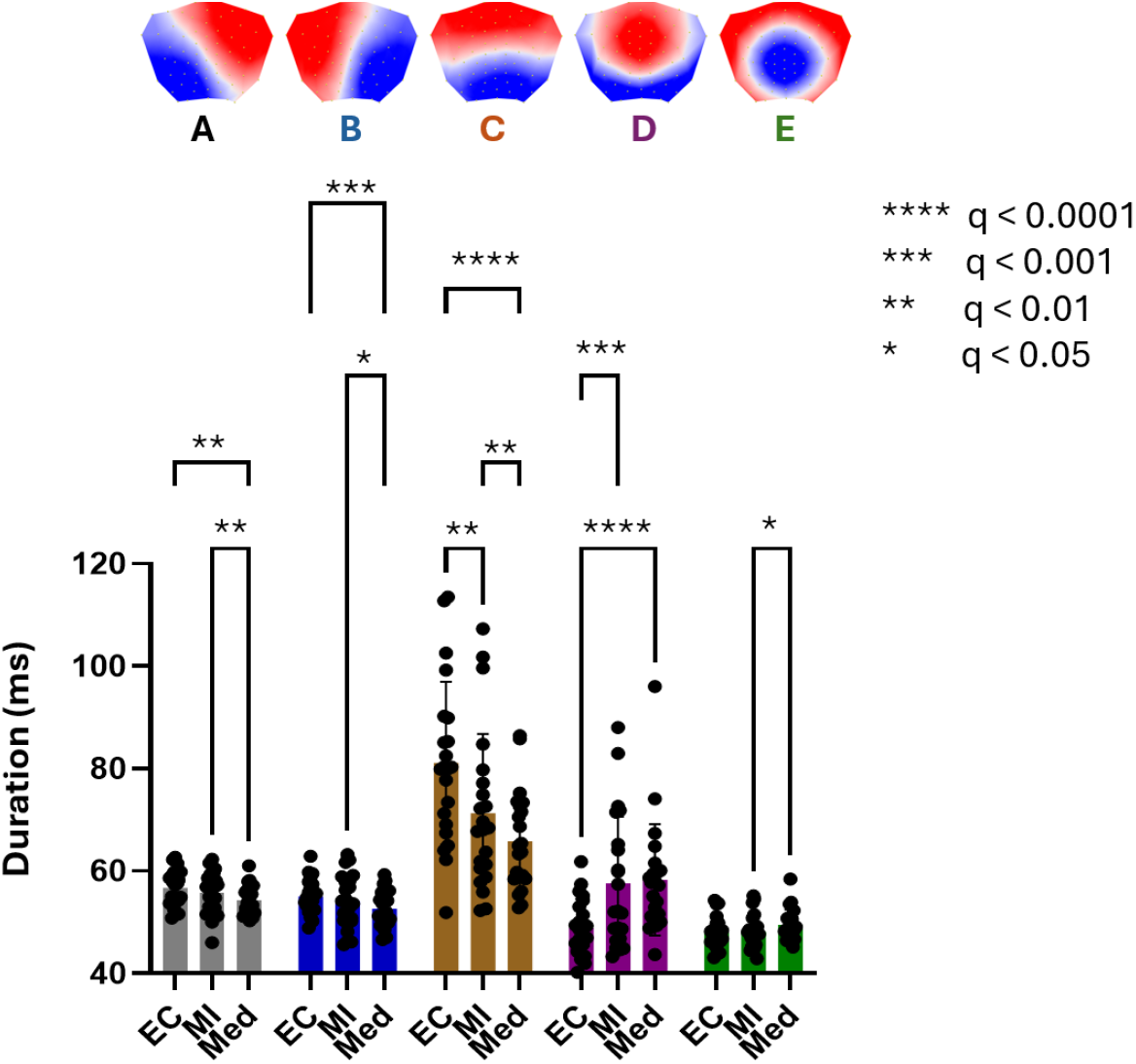
Microstate duration across conditions. Mean duration (ms) of microstates A–E across Baseline (BL), Mental Imagery (MI), and Meditation (Med). Microstate C showed longer overall durations than the other microstates. A significant Microstate × Condition interaction indicated reduced duration of microstates A–C during Meditation relative to BL and MI, increased duration of microstate D during Meditation relative to BL, and increased duration of microstate E during Meditation relative to MI. Mental Imagery relative to BL was associated with reduced duration of microstate C and increased duration of microstate E. Significance levels are indicated in the figure.

Post-hoc comparisons revealed that during Meditation compared to Baseline, microstates A, B, and C showed decreased duration, while microstate D showed increased duration. Compared to Mental Imagery, Meditation led to decreased duration of microstates A, B, and C, and increased duration of microstate E. Mental Imagery compared to Baseline resulted in decreased duration of microstate C and increased duration of microstate E.

### Occurrence

The two-way ANOVA revealed significant main effects for Microstate (F(1.852, 38.88) = 27.36; p < 0.0001) and Condition (F(1.612, 33.86) = 7.297; p < 0.01), as well as a significant Microstate × Condition interaction (F(3.745, 78.65) = 22.30; p < 0.0001). Post-hoc tests revealed that microstates D and E showed significantly increased occurrence during Meditation compared to both Baseline and Mental Imagery. Mental Imagery also showed significantly increased microstate D occurrence compared to Baseline (Figure 4).

**Figure 4:**
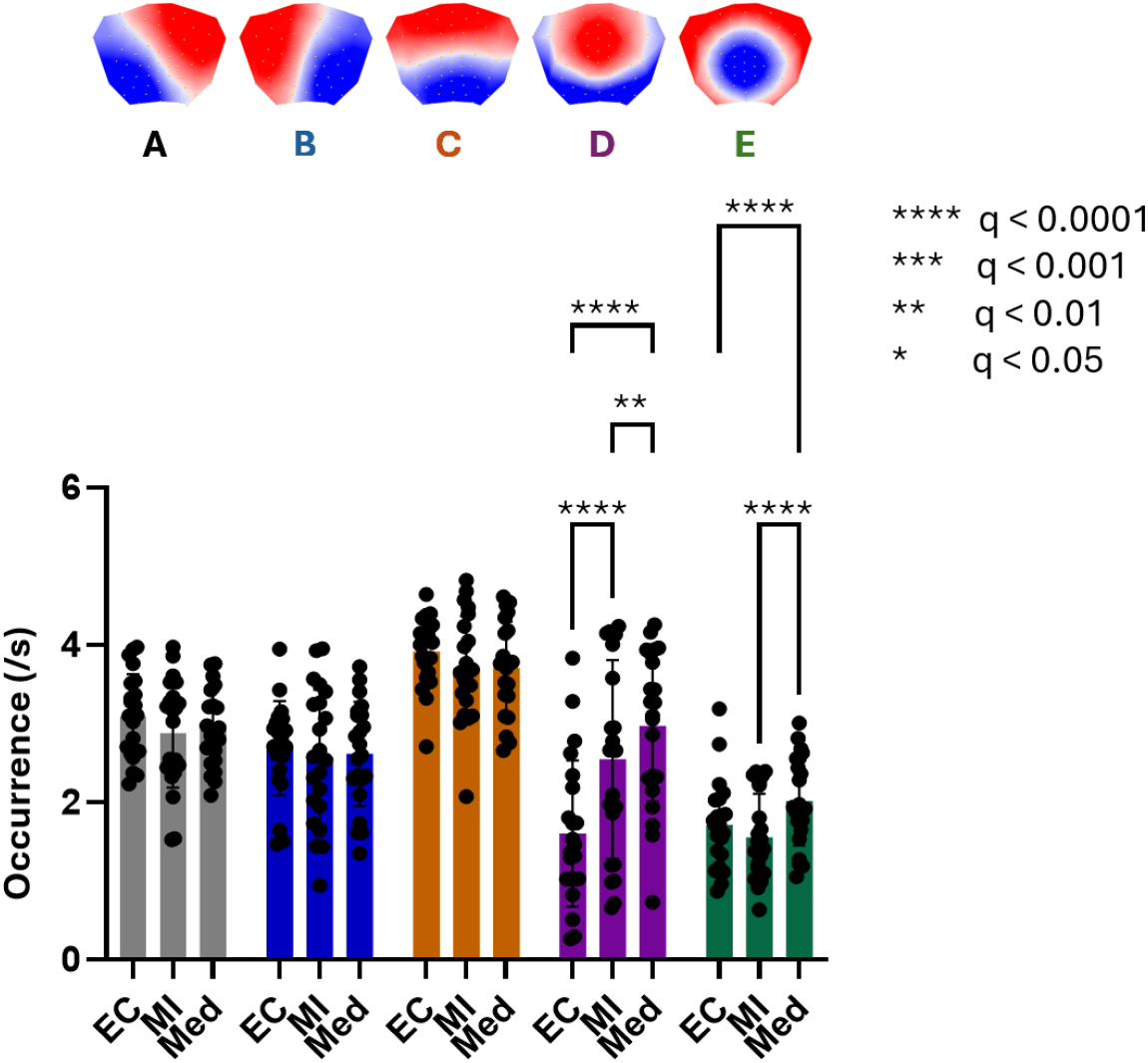
Microstate occurrence across conditions. Mean occurrence rate (events/s) of microstates A–E across Baseline (BL), Mental Imagery (MI), and Meditation (Med). A two-way repeated-measures ANOVA revealed significant main effects of Microstate and Condition, as well as a significant Microstate × Condition interaction. Post-hoc comparisons (FDR-corrected) showed increased occurrence of microstates D and E during Meditation relative to both BL and MI. Mental Imagery was also associated with increased occurrence of microstate D compared to BL. Significance levels are indicated in the figure.

### Source Localization

Source localization of the microstates revealed distinct networks for each microstate class (Figure 5 and Supplementary Figure 1). Microstates A and B showed activity in the right and left temporal pole, respectively. Microstate C, which decreased during meditation, was localized in the medial/lateral temporal cortex, encompassing the hippocampus, parahippocampal gyrus, and middle temporal gyrus. Microstate D, which increased during meditation, showed main activity in the posterior medial cortex, including the posterior cingulate cortex (PCC), precuneus, and cuneus bilaterally. Microstate E, which also increased during meditation, showed three main areas of activity: the left temporoparietal junction (TPJ) including the angular gyrus and supramarginal gyrus, the dorsolateral prefrontal cortex (DLPFC) bilaterally, and the left orbitolimbic region including the amygdala and extending to orbital frontal cortex.

**Figure 5:**
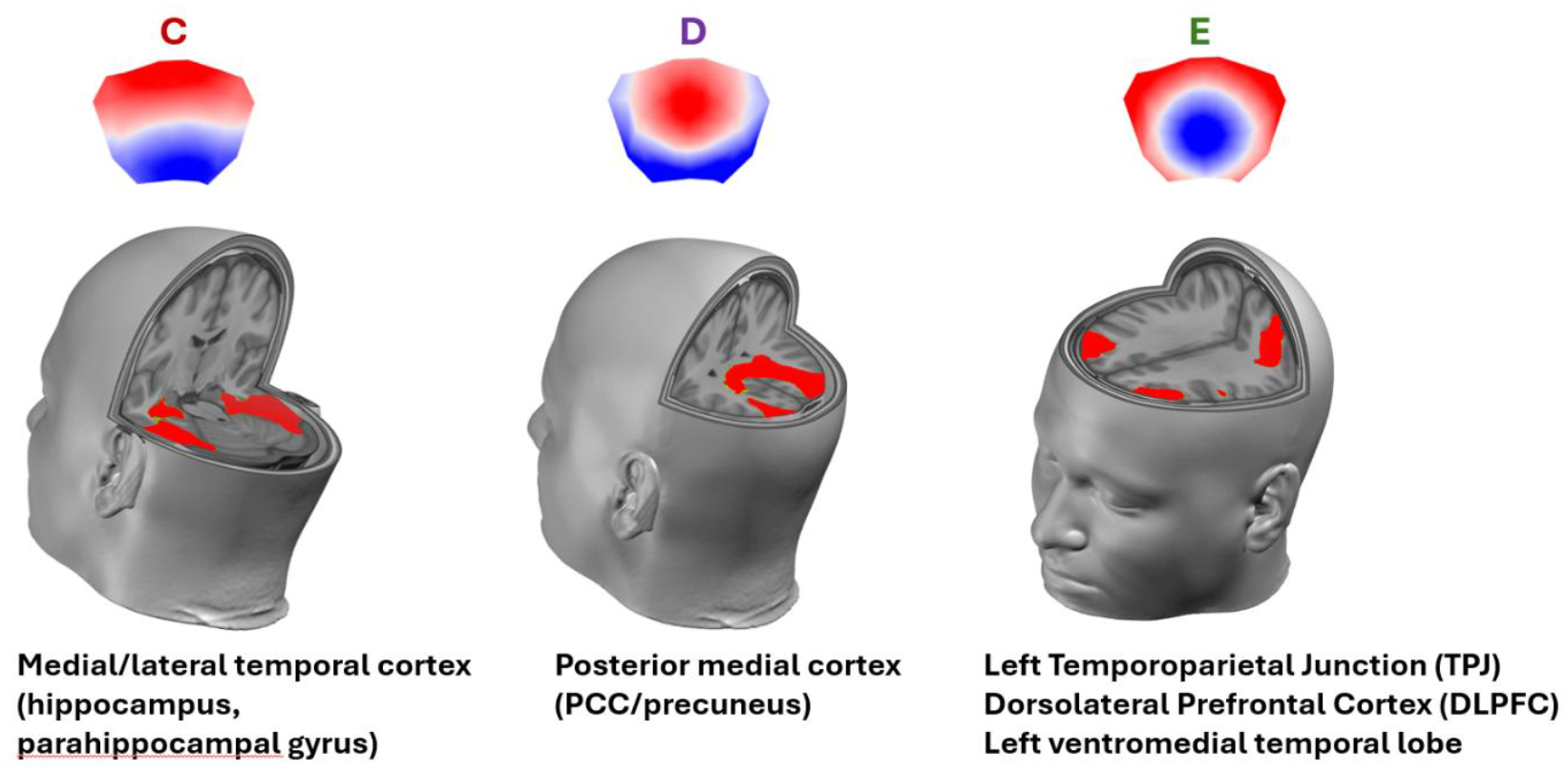
Distributed source estimates for microstates C–E, illustrating distinct cortical networks associated with each microstate class. Canonical scalp topographies are shown above the corresponding source maps.

**Supplementary Figure 1:**
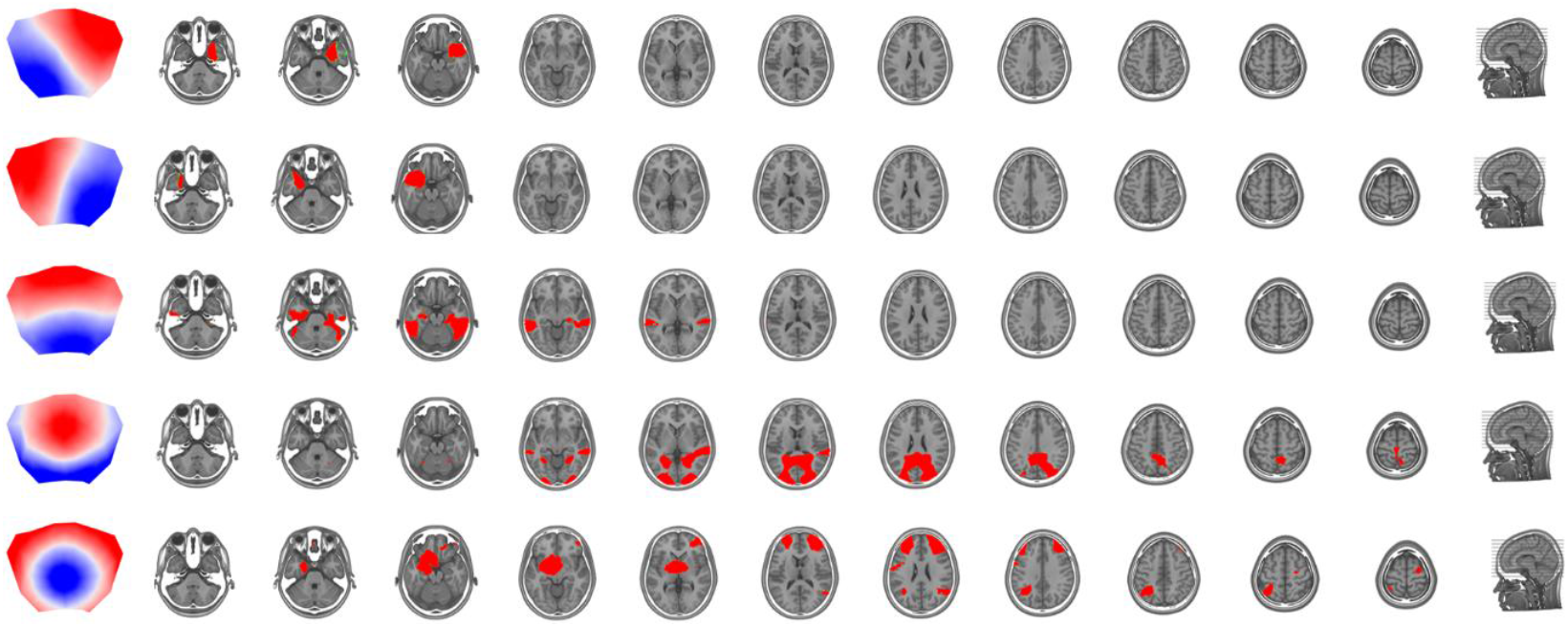
Whole-brain source distribution of EEG microstates. Slice-wise distributed source estimates for microstates A–E, shown across axial and sagittal planes on a standard anatomical template, with corresponding canonical scalp topographies.

## 4. Discussion

The present study investigated how focused-attention (FA) meditation on the breath modulates the temporal dynamics and neural generators of EEG microstates in experienced practitioners. Across multiple metrics—coverage, duration, and occurrence—we observed a consistent reorganization of microstate dynamics during meditation relative to both baseline rest and deliberate mental imagery. These findings extend the few prior work showing that meditation modulates the dynamics of large-scale neural networks measured with EEG microstates (Zarka et al., 2021; Faber et al., 2005, 2017; Panda et al., 2016).

Among all microstates, Microstate C emerged as the dominant temporal state during baseline, showing the longest mean duration as well as the highest occurrence and coverage. Relative to baseline, Microstate C was significantly reduced during both Mental Imagery and Meditation, with the strongest attenuation observed in the Meditation condition.

The effect of focused-attention (FA) meditation on Microstate C is comparable to that observed during non-autobiographical, externally oriented tasks such as mental arithmetic. Bréchet et al. (2019) reported reduced duration and occurrence of Microstate C during mental arithmetic compared with no-task rest and autobiographical memory tasks. Other studies have similarly shown longer Microstate C durations during unconstrained rest relative to arithmetic tasks (Kim et al., 2021; Seitzman et al., 2017). Croce et al. (2018), using magnetic stimulation, also linked Microstate C to task-negative mentation—mind-wandering, self-related thoughts, and emotional or interoceptive processing.

Source localization revealed that Microstate C was generated primarily within the hippocampus, parahippocampal gyrus, and middle temporal gyrus—regions strongly implicated in episodic memory retrieval and autobiographical processing during mind wandering (Faber & Mills, 2018; Christoff et al., 2016; Andrews-Hanna, 2012). This spatial profile aligns with prior evidence identifying Microstate C as being associated with autobiographical mentation (Tarailis et al., 2024) and are consistent with earlier work showing Microstate C sources in parietal and temporal brain regions, including the hippocampus (Custo et al., 2017; Bréchet et al., 2019; Bréchet et al., 2021; Tarailis et al., 2025). The suppression of Microstate C during meditation therefore likely reflects reduced engagement of hippocampal–temporal memory systems, a sub-network of the default mode network (DMN) (Andrews-Hanna, 2012), and a downregulation of spontaneous self-related thoughts and autobiographical mind-wandering. Within the framework of FA meditation, this reduction may index the practitioner’s capacity to detect the emergence of self-referential distractions and reorient attention toward the meditation object.

Microstates D and E showed complementary increases during meditation, reflecting a coordinated engagement of attentional control and self-regulatory systems. Microstate D, which increased in coverage, occurrence, and duration across both Mental Imagery and Meditation, was generated primarily in the posterior medial cortex, including the posterior cingulate cortex (PCC), precuneus, and cuneus, This network functions as a critical node for modulating attention between internal self-referential processing and external task demands (Leech & Sharp, 2014; Raichle et al., 2001). The PCC shows characteristic deactivation during externally focused, attentionally demanding tasks, while increased activity supports internally directed cognition (Leech et al., 2012), reflecting its role as a regulatory switch for attentional orientation (Leech et al., 2011).Previous studies have similarly localized Microstate D to parietal regions and have linked it to working memory, cognitive control, attentional reorientation, and the detection of behaviorally relevant stimuli (Croce et al., 2018; Tarailis et al., 2024). Increased presence of Microstate D has also been observed during arithmetic and recognition tasks compared with no-task rest (Bréchet et al., 2019; Kim et al., 2021; D’Croz-Baron et al., 2021; Murphy et al., 2018; Ferat et al., 2025). Additional work has associated greater Microstate D presence with heightened alertness to mental processing (Faber et al., 2017) and with transitions between internally and externally oriented cognition (Milz et al., 2016).

In the context of our study on focused-attention meditation, the increased presence of Microstate D suggests a mechanism by which the posterior medial cortex supports maintaining the internal task object (e.g., the breath at the nostril) and facilitates redirecting attention when distraction occurs. This pattern indicates that Microstate D reflects the establishment of a coherent internal attentional mode that becomes more prominent as participants transition from baseline into meditative states. The enhancement of Microstate D during meditation also aligns with fMRI findings demonstrating the PCC’s involvement in sustaining internal focus during meditative states (Kral et al., 2019).

Microstate E, which showed selective increases during Meditation, was localized to left temporoparietal junction (TPJ), the dorsolateral prefrontal cortex, and orbitolimbic areas including the amygdala. This three-component network provides a comprehensive neural substrate for meditative self-awareness. The TPJ is specifically implicated in self-other distinction processes and bodily self-awareness (Ionta et al., 2011; Blanke et al. 2015; Igelström and Graziano, 2017; Bréchet et al., 2018) critical for the sense of self during meditation. The DLPFC has been implicated in metacognitive self-awareness through both neuroimaging and brain stimulation studies (Fleming & Dolan, 2012; Lapate et al., 2020). Finally, the orbitolimbic region including amygdala and orbitofrontal cortex is associated with emotional self-awareness in meditation practitioners, with structural and functional changes in these regions correlating with mindfulness traits (Murakami et al., 2012; Baltruschat et al., 2021). This integrated network thus captures cognitive (DLPFC), perceptual (TPJ), and affective (orbitolimbic) dimensions of self-awareness characteristic of meditative states.

The findings also raise important questions for future research—particularly concerning the specific functional contributions of the DMN and the PCC during meditation. Although meditative training is often associated with reductions in DMN activity including the PCC (Brewer et al., 2011); Garrison et al. (2015) to diminish internal narrative processes and mind-wandering, recent work indicates that the DMN is not a uniform network but a constellation of interacting subsystems with potentially divergent roles. Prior studies have identified at least two major DMN subsystems—a medial temporal subsystem involved in memory-based simulation, and a dorsal medial subsystem linked to social-cognitive and conceptual processing—along with midline hubs such as the anterior medial prefrontal cortex and the PCC that integrate information across these components. These subsystems collectively support internally oriented cognition, including spontaneous thought, mental time travel, and reasoning about the self and others (Buckner et al., 2008; Andrews-Hanna et al., 2014). Our source localization results suggest that these DMN subsystems may be differentially reflected in distinct microstates: Microstate C corresponds to the medial temporal subsystem (hippocampus, parahippocampal cortex), Microstate D appears linked to PCC activity, and Microstate E aligns with regions associated with the dorsal medial subsystem, including the TPJ. This mapping implies that meditation may reorganize interactions *within* the DMN rather than simply suppressing it globally. This interpretation aligns well with the observation of Panda et al. (2016) in a simultaneous EEG-fMRI study where meditation decreased the connectivity of the PCC in the fMRI but increased the activity of the PCC-related microstate. This discrepancy highlights an important direction for future research: examining how the temporal properties of Microstate D relate to the stabilization of attention and determining whether its source localization remains anchored in the PCC across different attentional states. Such work could illuminate the nuanced interplay between DMN subsystems and attentional regulation during meditation.

## 5. Conclusion

Focused-attention meditation reliably reorganized EEG microstate dynamics. Compared with baseline rest and mental imagery, meditation produced a strong reduction of Microstate C (coverage, duration, occurrence) alongside increased Microstates D and E. Source localization suggests a shift away from medial/lateral temporal generators (hippocampus–parahippocampal regions; memory/self-generated mentation) toward posterior midline and frontoparietal/orbitolimbic networks supporting attentional stability, internal monitoring, and self-awareness. These results indicate that FA meditation reshapes the millisecond-scale temporal architecture of large-scale brain states.

## Notes

### Competing Interest Statement

Chuong Ngo, Monika Stasytyte, and Rodrigo Elizalde are employees of All Here SA. Erkin Bek is the founder of All Here SA. Christoph M. Michel is an independent contractor for All Here SA and provides EEG consultancy services through FBM-Analytics Sarl. Lionel Newman is an independent contractor for All Here SA and provides EEG computational services. The remaining authors declare no competing interests.

